# Gastrointestial absorption of pimozide is enhanced by inhibition of P-glycoprotein

**DOI:** 10.1101/2020.04.16.044602

**Authors:** Hiroki Morishita, Kozue Okawa, Misaki Ishii, Kenta Mizoi, Hiroshi Arakawa, Kentaro Yano, Takuo Ogihara

**Author notes:** These authors contributed equally to this work (Hiroki Morishita, Kozue Okawa, Kentaro Yano). These authors also contributed equally to this work (Misaki Ishii, Hiroshi Arakawa).

## Abstract

Following the death due to cardiac arrest of a patient taking pimozide, sertraline and aripiprazole antipsychotic/antidepressant combination therapy, a role of drug-drug interaction was suggested. Here, we investigated P-glycoprotein (P-gp)-mediated interaction among the three drugs using in vitro methods. Sertraline or aripiprazole significantly increased the permeability of pimozide in Caco-2 cell monolayers. ATPase assay indicated that pimozide is a P-gp substrate, and might act as a P-gp inhibitor at higher concentrations. The values of the kinetic parameters of carrier-mediated efflux, calculated from the concentration dependence of pimozide efflux from LLC-GA5-COL150 cells expressing human P-gp, were as follows: maximum transport rate (J_max_) = 84.9 ± 8.9 pmol/min/mg protein, half-saturation concentration (K_t_) = 10.6 ± 4.7 μM, first-order rate constant (k_d_) = 0.67 ± 0.14 pmol/min/mg protein. Further, the efflux ratio of pimozide in LLC-GA5-COL150 cells was significantly decreased in the presence of sertraline or aripiprazole. These results indicate that pimozide is a substrate of P-gp, and its efflux is inhibited by sertraline and aripiprazole. Thus, P-gp inhibition by sertraline and/or aripiprazole may alter the gastrointestinal permeability of co-administered pimozide, resulting in an increased blood concentration of pimozide, which may increase the likelihood of pimozide’s known life-threatening side effect of QT prolongation.

## Introduction

Pimozide is an antipsychotic used to treat schizophrenic and pediatric autistic disorders. However, one of its major side effects is QT prolongation [1], because it is a strong antagonist of the alpha subunit of a potassium ion channel (hERG) [2] and this action causes a significant prolongation of QT intervals [3]. Several antidepressants are restricted for use in combination with pimozide, because of the increased risk of this side effect [4]. For example, the combination of pimozide and sertraline is contraindicated because it leads to an increased blood concentration of pimozide, thus increasing the risk of QT prolongation. Nevertheless, a case has been reported in which a male child administered pimozide together with sertraline and aripiprazole died due to cardiac arrest [5]. Pimozide is a substrate of the metabolic enzymes CYP3A4, 2D6, and 1A2, and aripiprazole is a substrate of 3A4 and 2D6 [6], while sertraline is a substrate of 2C19, 2C9, 2B6, and 3A4, and a mild to moderate inhibitor of CYP2D6 [7]. Therefore, it had been suspected that interaction among those drugs would be due to inhibition of CYP-mediated pimozide metabolism by sertraline and/or aripiprazole. On the other hand, Alderman reported that the C_max_ and AUC of pimozide were increased by 35% and 37%, respectively, with no significant difference in blood half-life, when the drug was used in combination with sertraline [8]. In pharmacokinetics, blood half-life is inversely proportional to clearance; in other words, if hepatic CYP3A4 and/or CYP2D6 metabolism were inhibited by sertraline, the blood half-life of pimozide should be prolonged. This suggests that a drug interaction mechanism(s) other than inhibition of CYP3A4 and/or CYP2D6 is involved. We speculated that the drug transporter P-glycoprotein (P-gp) might be an interaction site.

P-gp is a member of the ATP-binding cassette superfamily, and is mainly localized at the intestine, blood-brain barrier, adrenal gland, enterocytes, hepatocytes, placenta and renal proximal tubules in humans [9]. P-gp is responsible for the efflux of many xenobiotics and plays major roles in drug absorption, distribution and excretion. In the intestine, P-gp mediates the efflux of its substrates, restricting the absorption of many xenobiotics, including drugs [10–13]. Recent studies show that sertraline, aripiprazole, and several of their metabolites have a P-gp-inhibitory effect [14–16], whereas pimozide is not known to be a P-gp substrate. The bioavailability of pimozide is limited (about 50%) [7]. Therefore, if pimozide is a P-gp substrate, there is a possibility that its absorption could be increased when it is used in combination with P-gp inhibitors. In order to test this idea, we examined whether pimozide is a P-gp substrate, and whether its gastrointestinal permeability might be influenced by sertraline and/or aripiprazole by means of a series of in vitro studies.

## Materials and Methods

### Chemicals

Pimozide was purchased from Santa Cruz Biotechnology (Dallas, TX). Sertraline hydrochloide, aripiprazole and verapamil hydrochloride were purchased from Wako Pure Chemical Industries (Osaka, Japan). All other reagents were commercial products of reagent grade.

### Cell culture

Caco-2 cells, a human colon epithelial cancer cell line frequently used as an in vitro intestinal model, were purchased from the American Type Culture Collection (Rockville, MD, USA). They were cultured, passaged and grown as described previously [19] in Dulbecco’s modified Eagle’s medium supplemented with 10% fetal bovine serum, 100 units/ml penicillin, 0.1 mg/ml streptomycin, and 1.0 % nonessential amino acids, at 37°C in an atmosphere of 5% CO_2_ in air. The cells were seeded on Transwell filter membrane inserts (Costar, Bedford, MA, USA) at a density of 6 ×10^4^ cells/cm^2^. The culture medium was replaced with fresh medium every second or third day and Caco-2 cell monolayers grown for 21 days were used for the transport studies. Transepithelial electrical resistance (TEER) was measured using a Millicell-ERS resistance system (Millipore, Bedford, MA, USA). Cell monolayers used for transport studies had TEER values of 800 to 1000 Ω·cm^2^.

LLC-GA5-COL150 cells, a derivative of kidney epithelial cell line LLC-PK1 expressing human P-gp on the apical membrane, were obtained from Riken Gene Bank (Tsukuba, Japan). They were cultured, passaged and grown as described previously [20] in Medium 199 supplemented with 10% fetal bovine serum, 100 U/mL penicillin, 100 µg/mL streptomycin and 150 ng/mL colchicine, at 37°C in an atmosphere of 5% CO_2_ in air. The cells were seeded on Transwell filter membrane inserts (Costar, Bedford, MA, USA) at a density of 2.5 × 10^5^ cells/cm^2^ for transport studies [20], and seeded on 24-well cell culture plates (Corning, NY, USA) at a density of 1.5 × 10^5^ cells/cm^2^ for efflux assays [21]. The culture medium was replaced with fresh medium every second or third day. Cells were grown for 7 days and used for experiments. One day prior to experiments, the medium was changed to Medium 199 without colchicine. Cell monolayers with TEER values of 300 to 600 Ω·cm^2^ were used for transport studies.

### Transport experiments

In the transport studies with Caco-2 cell monolayers, the transport medium on the apical side consisted of Hanks’ balanced salt solution (HBSS) with 10 mM 2-morpholinoethanesulfonic acid (MES) (pH 6.5) and that on the basal side consisted of HBSS with 10 mM 2-[4-(2-hydroxyethyl)-1-piperazinyl]ethanesulfonic acid (HEPES) (pH 7.4). Test chemicals were dissolved in dimethyl sulfoxide (DMSO) and diluted with transport medium (the final DMSO concentration was 1%). Caco-2 cell monolayers were preincubated with the transport medium for 20 min at 37ºC. Transport experiments were initiated by adding the transport medium containing pimozide (10 μM) to the donor side, while the receiver side was filled with transport medium. The concentration of pimozide was chosen with reference to the clinical dose of 1 mg dissolved in 250 mL of water [22], which corresponds to a concentration of 9 μM. Inhibitor was added to both sides during inhibition studies. Sertraline (500 μM), aripiprazole (15 μM) and verapamil were used as inhibitors (verapamil as a positive control). The tested concentrations of sertraline and aripiprazole were chosen in the same manner as described above, based on the calculation that the clinical doses of sertraline (25 mg) and aripiprazole (3 mg) in 250 mL of water would correspond to concentrations of 292 μM and 27 μM, respectively. Samples were collected at 20, 40, 60, 90 and 120 min, and replaced with equal amounts of transport medium. In the transport studies with LLC-GA5-COL150 cell monolayers, the transport medium on both sides consisted of HBSS with 10 mM HEPES (pH 7.4). Inhibitors were added to the donor side and transport experiments were initiated in the same manner. Samples were collected at 15, 30, 45 and 60 min, and replaced with equal amounts of transport medium. The time course of drug transport in the apical (A) to basal (B) direction (A to B) and that in the opposite direction (B to A) were observed at 37°C. The permeability (P_app_) across cell monolayers was evaluated by dividing the slope of the experimental time course in the A-to-B or B-to-A direction by the concentration on the donor side and is represented as P_app AtoB_ or P_app BtoA_, respectively. The efflux ratio (ER) was calculated by dividing P_app BtoA_ by P_app AtoB_.

### ATPase assay

The SB-MDR1 PREDEASYTM ATPase Kit, including P-gp-expressing membrane vesicles, was purchased from SOLVO Biotechnology (distributed by KAC Co., Ltd., Kyoto, Japan). This assay kit is designed to measure the interaction of test drugs with P-gp. ATPase assay was conducted according to the manufacturer’s instructions and a previous report [18]. Inorganic phosphate liberated by ATP hydrolysis was detected by colorimetric reaction. The optical density (OD) was measured at 590 nm with a microplate reader, Sunrise™ Rainbow (TECAN, Kanagawa, Japan).

### Efflux study

To investigate the involvement of P-gp in pimozide efflux transport, efflux studies with LLC-GA5-COL150 cells were performed. Cells seeded on 24-well cell culture plates were washed twice with ice-cold Dulbecco’s phosphate-buffered saline (-) (D-PBS(-)), and pretreated with 300 µL of ice-cold Opti-MEM containing 0.01 to 100 µM pimozide for 30 min on ice (4℃). The Opti-MEM was removed, and the cells were washed twice with ice-cold D-PBS(-). Experiments were initiated by adding pimozide-free Opti-MEM to each well and incubating the plates at 37℃. After 10 min, transport was stopped by washing each well three times with ice-cold D-PBS(-), and the cells were lysed by adding 200 µL of 0.1 N NaOH solution. Cell lysates were used for protein determination and measurement of pimozide concentration. Protein was determined colorimetrically using DCTM Protein Assay (BIO-RAD, Hercules, CA), based on absorbance measurement at 700 nm with a microplate reader, Sunrise Rainbow RC (Tecan, Kanagawa, Japan). An aliquot of cell lysate was mixed with 300 µL of ethyl acetate for 10 min in the cold, then 100 µL of the organic phase was moved to a new tube and evaporated. The residue was dissolved in 300 µL of mobile phase. All samples were applied to MultiScreen® Solvent Filter Plates 0.45 µm Low-Binding Hydrophilic PTFE (Merck, Ireland) and centrifuged at 3,500 rpm for 20 min at 4℃. Filtered samples were collected in a 96-well microplate (Asone, Japan), and pimozide concentration was determined by triple quadrupole liquid chromatography mass spectrometry (LC-MS/MS) as described below. Efflux rate was calculated according to the following equation.

Efflux rate = (X_0_ – X_1_) / incubation time

Where X_0_ is pimozide concentration per protein amount before incubation and X_1_ is that after incubation. Therefore, X_0_ – X_1_ is the net transport of pimozide from LLC-GA5-COL150 cells.

### Kinetic analysis

To estimate the kinetic parameters of carrier-mediated transport in the efflux assays, the transport rate (J) was fitted to the following equation (1), containing saturable and nonsaturable-linear terms, by using the nonlinear least-square regression analysis program, MULTI, as previously reported [23].

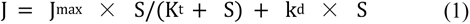

Where J_max_ is the maximum transport rate for the carrier-mediated transport, S is the substrate concentration, K_t_ is the half-saturation concentration, and k_d_ is the first-order rate constant.

### Measurement of pimozide

Concentrations of pimozide were determined by LC-MS/MS analysis. Pimozide samples (10 µL) were injected into an HPLC system (LC-20A system, Shimadzu, Kyoto, Japan) equipped with a CAPCELL PAK C18 MGⅢ / 3 µm column (φ2.0 × 50 mm, Shiseido Co. Ltd., Tokyo, Japan). The mobile phase consisted of a mixture of acetonitrile containing 0.1% formic acid (organic solvent phase) and distilled water containing 0.1% formic acid (water phase) (50: 50). The flow rate was 0.1 mL/min, at 40℃. Analytes were detected using a quadrupole mass spectrometer (LCMS-2010EV, Shimadzu) fitted with an electrospray ionization source. Analytes were detected in the positive mode, and the protonated molecular ion of pimozide was monitored at *m*/*z =* 109.05.

### Statistical analysis

Kinetic parameters are presented as mean ± standard deviation (S.D.). Other data are presented as mean ± standard error of the mean (S.E.M.). Statistical analysis of kinetic parameters was performed by means of Dunnet’s *t-*test. A difference between means was considered to be significant when the *P* value was less than 0.05.

## Results

### Transport study across Caco-2 cell monolayers

The P_app_ of pimozide in the A-to-B direction in the presence of sertraline was 5.9-fold higher than that in the absence of sertraline. The P_app_ in the B-to-A direction was decreased by sertraline, aripiprazole and verapamil, and sertraline had the greatest effect (Fig 1A). Overall, the addition of sertraline, aripiprazole and verapamil decreased the efflux ratio to 2.9 %, 60.6 % and 43.9 % respectively, compared with pimozide alone (Fig 1B).

**Fig 1.**
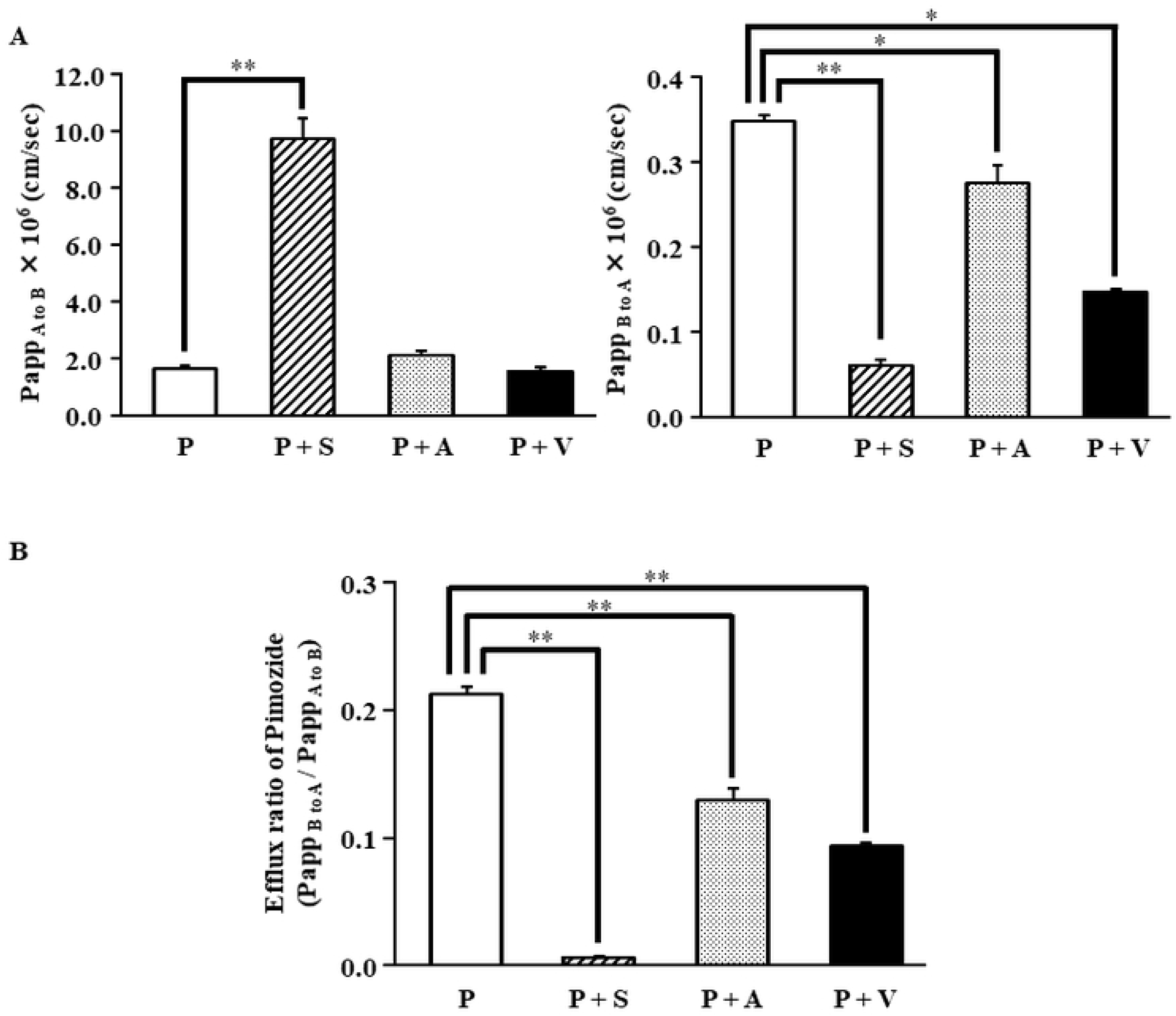
Effects of sertraline, aripiprazole and verapamil on pimozide transport across Caco-2 cell monolayers. The permeability of pimozide at a concentration of 10 µM was measured in Caco-2 cell monolayers in the absence or presence of sertraline, aripiprazole or verapamil. Bars represent P_app_ of pimozide under each condition (Fig 1A). The efflux ratio of pimozide was calculated from P_app_ under each condition (Fig 1B). Data are shown as mean ± SEM (n = 3). P: Pimozide 10 µM. S: Sertraline 500 µM. A: Aripiprazole 15 µM. V: Verapamil 100 µM. **, *p* < 0.01 compared with pimozide.

### ATPase assay for P-gp substrate

We investigated the pimozide concentration-sensitive ATPase activity of P-gp (Fig 2). Concentration-dependent elevation of ATPase activity was initially observed, though concentrations of pimozide over about 10 µM decreased the ATPase activity.

**Fig 2.**
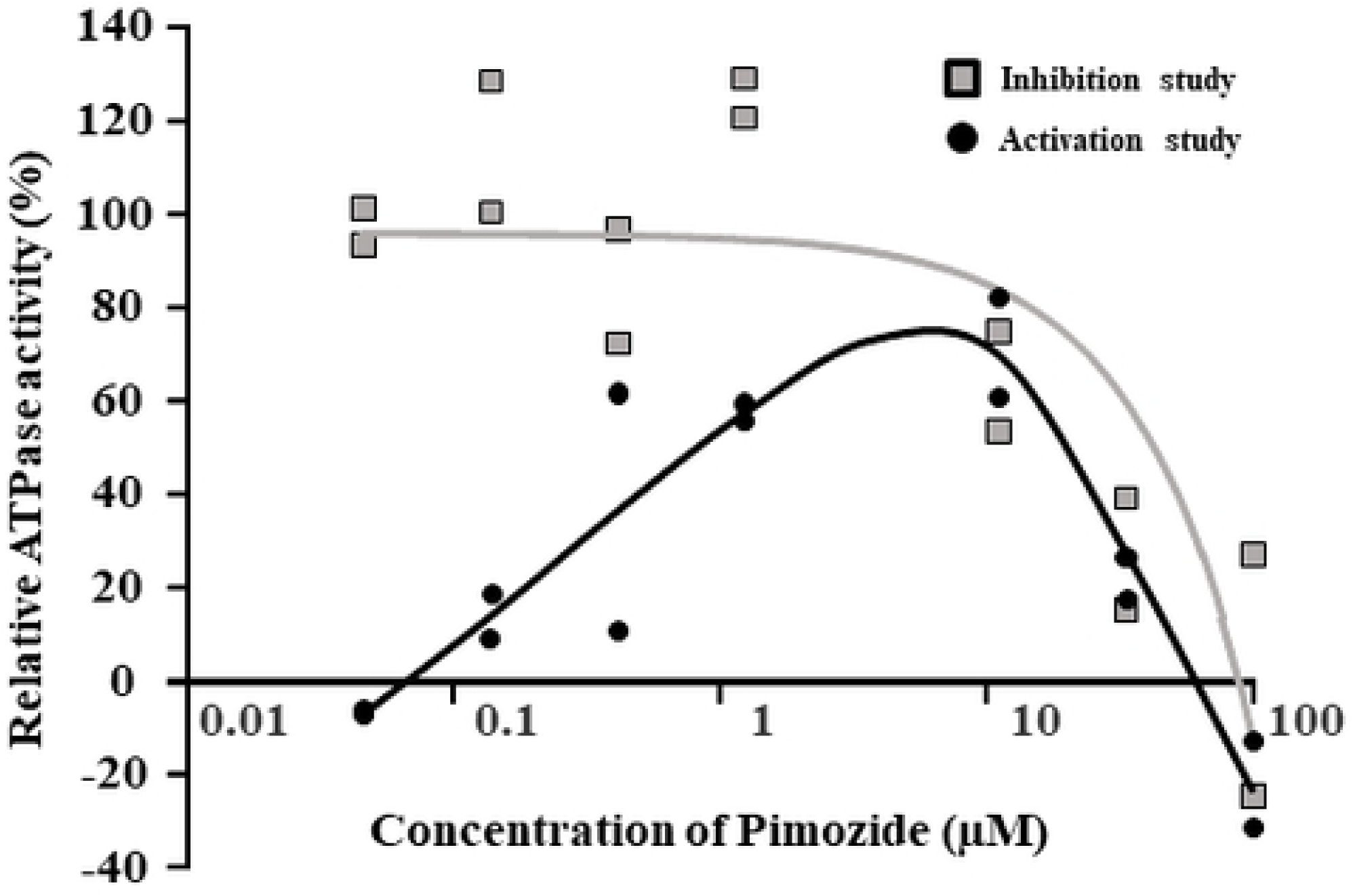
ATPase activity of pimozide. The results of ATPase activity assay are shown. Solid circles show activation study (n = 2) and open squares show inhibition study (n = 2). The black line represents the average and the gray line is the fitted curve. Data are given as relative activity (% of control). The pimozide concentration range is from 0.04 µM to 100 µM.

### Efflux study of pimozide

The concentration dependence (in the range of 0.01 to 100 µM) of pimozide efflux from LLC-GA5-COL150 cells is shown in Fig 3. The J_max_, K_t_ and k_d_ values estimated from the efflux assay data according to the equation (1) given in Materials and Methods were 84.9 ± 8.9 pmol/min/mg protein, 10.6 ± 4.7 μM and 0.67 ± 0.14 pmol/min/mg protein, respectively.

**Fig 3.**
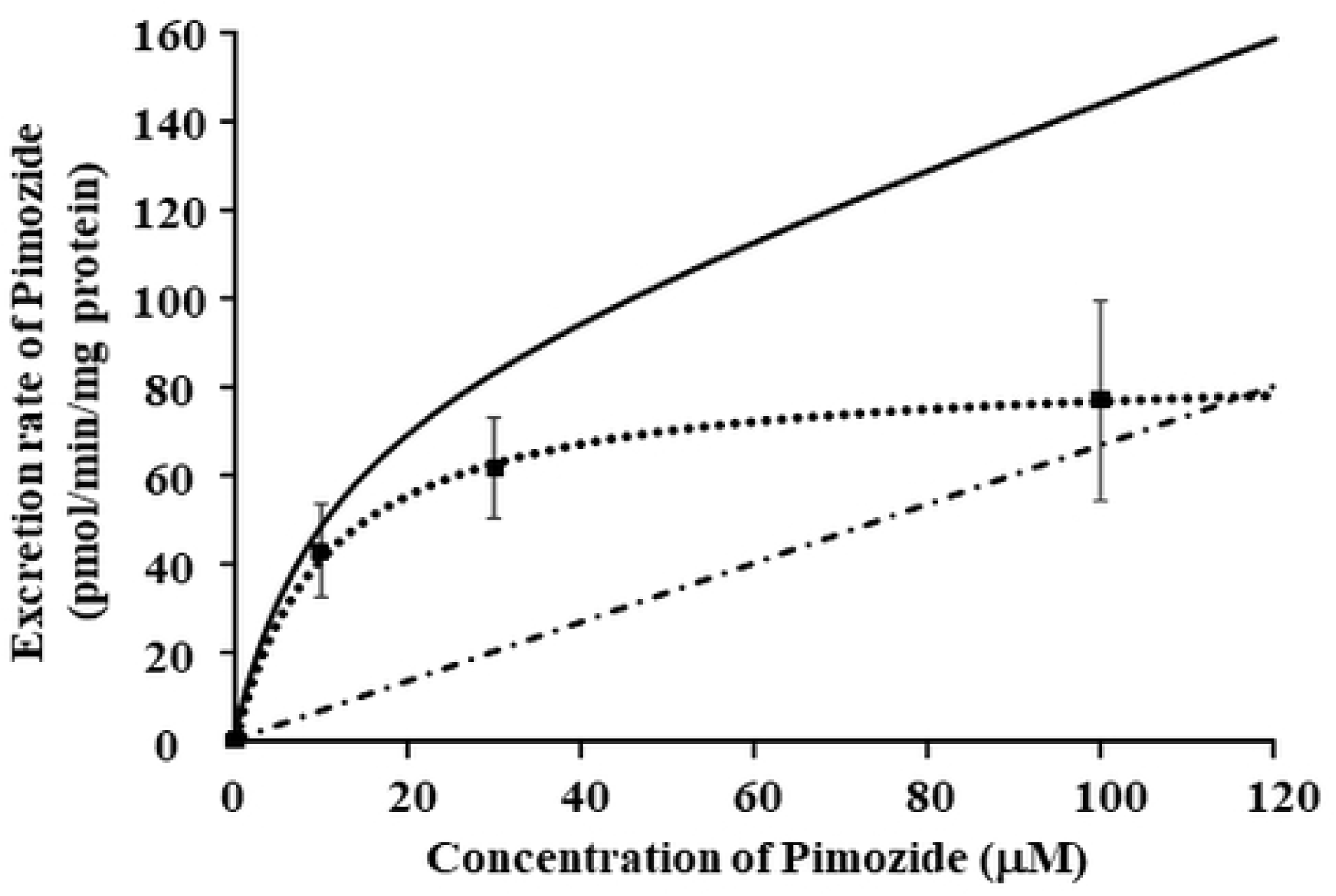
P-gp-mediated efflux transport of pimozide in LLC-GA5-COL150 cells. The efflux rate of pimozide from LLC-GA5-COL150 cells are shown. The black line represents the efflux rate of pimozide from LLC-GA5-COL150 cells. The broken lines represent the saturable (••••) and non-saturable (-•-•-) transport components. The pimozide concentration range is from 0.01 µM to 100 µM.

### Transport study across P-gp-expressing LLC-GA5-COL150 cell monolayers

Sertraline and aripiprazole significantly decreased the efflux ratio in LLC-GA5-COL150 cell monolayers to 11.0 and 21.9 % of the control, respectively, compared with pimozide alone (Fig 4).

**Fig 4.**
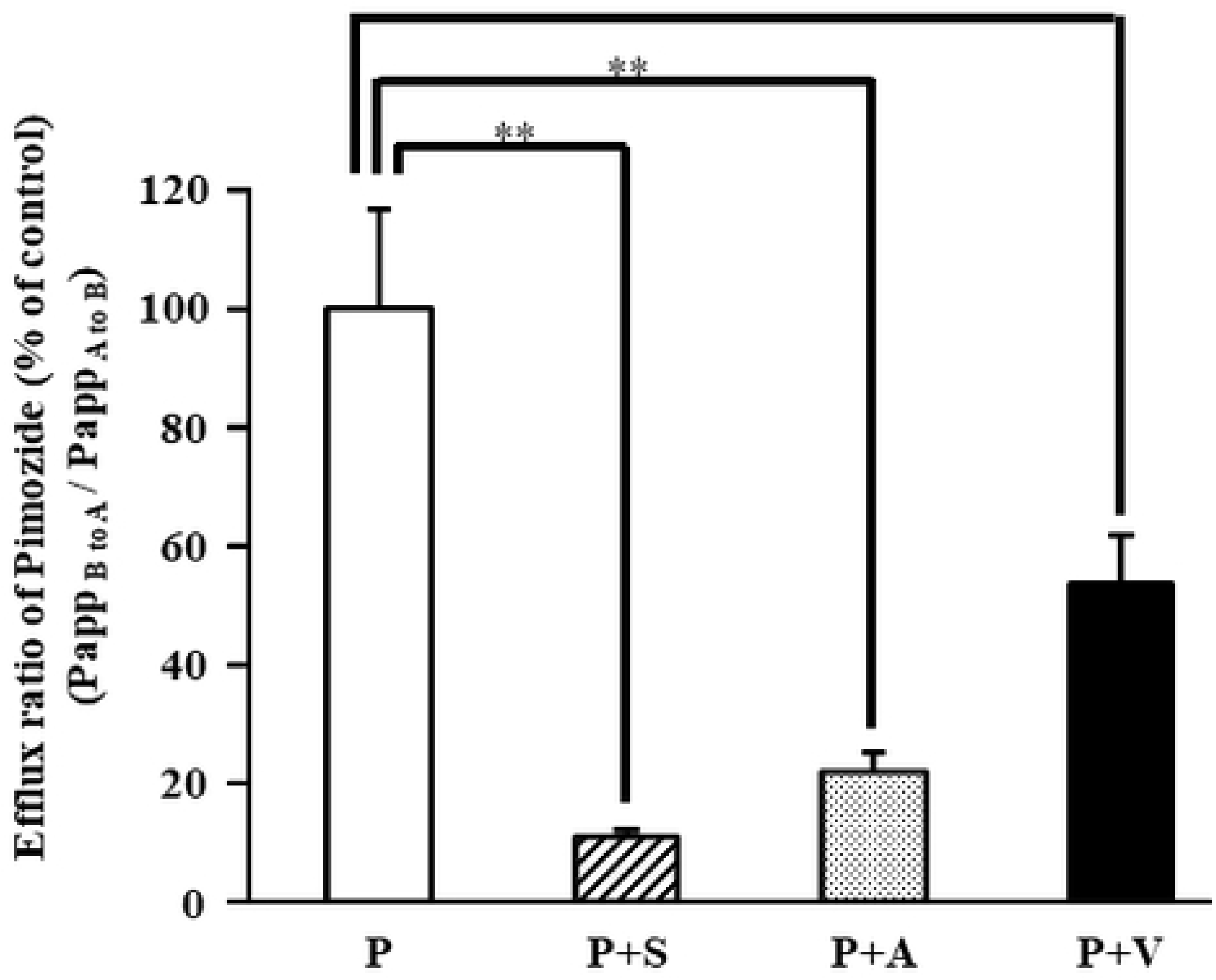
Efflux ratio of pimozide in LLC-GA5-COL150 cell monolayers. The values of the efflux ratio of pimozide in the absence or presence of sertraline, aripiprazole or verapamil in LLC-GA5-COL150 cell monolayers are shown. Data are shown as mean ± SEM (n = 5~6). Values were normalized to the control. P: Pimozide 10 µM. S: Sertraline 500 µM. A: Aripiprazole 15 µM. V: Verapamil 100 µM. *, *p* < 0.05; **, *p* < 0.01 compared with pimozide.

## Discussion

To evaluate the possibility of drug-drug interaction of pimozide with sertraline and/or aripiprazole in the gastrointestinal tract, we performed a transport study with Caco-2 cells, which have been widely used for experimental studies of gastrointestinal permeability. The ER of pimozide was decreased significantly and the permeability was increased when sertraline or aripiprazole was co-administered with pimozide, compared with the ER of pimozide alone (Fig 1B). Moreover, the ATPase activity of P-gp-expressing inverted membrane vesicles was elevated in a pimozide-concentration-dependent manner (Fig 2). However, the ATPase activity was reduced in the high concentration range, suggesting that pimozide is both a P-gp substrate and an inhibitor at higher concentrations. Futhermore, the efflux of pimozide from P-gpoverexpressing LLC-GA5-COL150 cells had both saturable and non-saturable linear components (Fig 3). For the saturable carrier-mediated transport of pimozide, we obtained J_max_ and K_t_ values of 84.9 pmol/min/mg protein and 10.6 µM, respectively. Therefore, J_max_/K_t_, which reflects affinity for the transport carrier(s), is 8.0 µL/min/mg protein. J_max_ and K_t_ of Rhodamine123 (Rho123), a well-known P-gp substrate, are 988 pmol/min/mg protein and 173 µM, respectively [24], giving a J_max_/K_t_ value of 5.7 µL/min/mg protein. Therefore, the affinity of pimozide for P-gp may be similar to that of Rho123, although it should be noted that the experimental method and conditions were not the same in the two cases. Pimozide was transported in the B-to-A direction of LLC-GA5-COL150 cell monolayers, and sertraline and aripiprazole decreased the ER of pimozide by about 90% and 80%, respectively (Fig 4). In the gastrointestinal tract, the effective concentration of pimozide is estimated to be at least 9 μM and its K_t_ for P-gp was 10.6 μM, suggesting that P-gp-mediated drug interactions could occur. Importantly, because the experimental concentrations of sertraline and aripiprazole were set based upon the clinically used doses, it is likely that P-gp-mediated drug interaction in the gastrointestinal tract could occur in clinical situations when these drugs are co-administered with pimozide.

In conclusion, pimozide is a substrate of P-gp, and its absorption is increased by sertraline and aripiprazole. Our results suggest that elevated pimozide blood levels observed when the drug is administered in combination with sertraline and/or aripiprazole can explained at least in part by interaction at P-gp. Further in vivo studies seem warranted.

## Acknowledgments

This research was supported by the Laboratory of Biopharmaceutics, Department of Phamacology, Faculty of Pharmacy, Takasaki University of Health and Walfare, and by the Gunma Yakugakunetwork.

## Author contributions

Investigation: Hiroki Morishita, Kozue Okawa, Misaki Ishii, Kenta Mizoi, Hiroshi Arakawa, Kentaro Yano.

Methodology: Hiroki Morishita, Kozue Okawa, Misaki Ishii. Project administration: Hiroki Morishita, Kentaro Yano.

Supervision: Takuo Ogihara

Writing – original draft: Hiroki Morishita Writing – review & editing: Takuo Ogihara

